# Ultra-small bacteria and their phages discovered through metagenomics in a sub-arctic degrading permafrost ecosystem

**DOI:** 10.64898/2026.11.11.765320

**Authors:** Nea Tuomela, Tatiana Demina, David N. Thomas, Hermanni Kaartokallio, Jenni Hultman

## Abstract

Patescibacteriota is a diverse bacterial phylum characterized by ultra-small cell size, streamlined genome, incomplete metabolism and episymbiotic lifestyle. While Patescibacteriota have been detected across a wide range of ecosystem types, their phages are not widely described. We utilized metagenomics to explore these less characterized parts of the microbiome in a sporadic permafrost ecosystem, with samples from palsa and peat soils, small ponds and streams. We retrieved 33 high-quality metagenome assembled genomes (MAGs) of Patescibacteriota, mainly from the classes Minisyncoccia and Saccharimonadia. Minisyncoccia was dominant class in the thawing soils and thaw ponds, whereas Saccharimonadia was more abundant in the frozen palsa soil layers. The distribution of the Patescibacteriota classes was reflected in their metabolic potential. For example, gluconeogenesis pathway was complete only in MAGs from the class Minisyncoccia, whereas cytochrome o ubiquinol oxidase was mainly found from the class Saccharimondia. The viral community analysis yielded 16 354 viral operational taxonomic units (vOTUs), which represented mainly tailed dsDNA viruses from the class *Caudoviricetes*. Based on genome size, 16 vOTUs were qualified as jumbophages and one as a megaphage. Similar to the bacterial community, the viral community composition differed across sample types. The most common hosts for bacterial viruses were phyla Pseudomonadota, Acidobacteriota and Actinomycetota. A further 196 vOTUs were *in silico* linked to Patescibacteriota hosts. The Patescibacteriota phages were diverse, with only two vOTUs representing the same genus. The relative abundance of Patescibacteriota-associated phages was especially high in frozen palsa soils, largely following the same pattern of abundance as Patescibacteriota.

## Introduction

Peatland and permafrost soils are two of the largest terrestrial carbon reservoirs facing rapid changes as temperatures in the Arctic region increase [1,2]. Permafrost thaw is accelerated by climate change [3], leading to irreversible changes in carbon storage through mainly increased microbial activity [4,5]. In addition to soil, small-scale freshwaters like ponds and streams can also act as hotspots for greenhouse gas emissions (GHG) [6,7].

Patescibacteriota (previously Candidate Phyla Radiation) is an enigmatic phylum representing a significant proportion of previously unknown bacterial diversity [8,9]. The abundance of Patescibacteriota has been likely underestimated with standard sampling methods, especially in aquatic ecosystems. The standardly used filter pore size of 0.22 µm does not capture all Patescibacteriota due to their small cell size [10,11], and 16S rRNA gene sequencing fails to capture all Patescibacteriota due to primer bias [12]. However, Patescibacteriota have been established as one of the most abundant phyla in a variety of ecosystems, including groundwater [13–15], permafrost thaw ponds [16], permafrost soils [17] and wastewaters [18,19].

In addition to ultra-small cell size, Patescibacteriota are characterized by streamlined genomes and incomplete basic metabolic pathways [20–22]. The lack of essential pathways and reduced genomes are complemented through obligatory episymbiotic and parasitic lifestyles [23–26]. Patescibacteriota-host pairings have been studied through a few cultured strains; e.g. representative of Saccharimonadia has been linked to Actinomycetota host [25], Minisyncoccia to methanogenic archaea [24,27] and Gracilibacteria to methylotrophic Pseudomonadota [28]. Patescibacteriota has been shown to impact the hosts’ cell morphology [29] and population growth by slowing the cell division of infected host cells [25]. Patescibacteriota might potentially have population level impact, and with methanogenic or methanotrophic host, Patescibacteriota parasitism could further impact GHG dynamics.

Viral communities are thought to have significant roles in biogeochemical cycles including carbon turnover in permafrost soils [30]. Viruses can cause the release of nutrients and carbon through lysis of their host cells or potentially enhance the metabolic capacity of their hosts through auxiliary metabolic genes (AMGs) [31]. In permafrost ecosystems, viruses can support their hosts’ ability to degrade complex and labile organic compounds [30,32]. Patescibacteriota are predicted to be hosts to diverse viruses, creating a complex interaction involving phages infecting Patescibacteriota and Patescibacteriota depending on other bacterial or archaeal hosts [28,33,34]. Until now, Patescibacteriota phages have mainly been predicted through metagenomics in groundwater [28,33,35] and human microbiome [34]. Recently, a phage induced from the bacterial host of a Patescibacteriota epibiont was found to also infect the epibiont, providing the first experimental evidence of a tripartite phage-host-Patescibacteriota interaction [36].

The aim in this study was to characterize taxonomy, abundance and metabolic potential of Patescibacteriota in a sub-arctic permafrost palsa ecosystem. In addition, we explored the viral community composition across the permafrost peatland ecosystem. We linked viruses to their putative hosts, including Patescibacteriota, and explored their lifestyles and functional potential in this climate-wise vulnerable ecosystem.

## Materials and methods

### Site description and sampling

Samples and the dataset were utilized from previously collected soil profiles of six palsa and five peat soil cores, their adjacent ponds and stream water samples from Vaisjäeggi palsa mire in Kevo, Finland [37]. Water samples were sequentially filtered up to 0.1µm with PES membrane filters, and DNA was extracted from the filters and bulk soil with PowerSoil Pro Kit. The libraries were prepped with Nextera XP at the DNA Sequencing core lab at University of Helsinki and DNA sequenced with AVITI [37]. Metagenomes were quality filtered, assembled and binned for MAG retrieval as described elsewhere [38], and viral analysis was done on the bulk metagenomes. MAGs were utilized in the analysis for viral host predictions, and Patescibacteriota MAGs were further explored for their metabolic potential and abundance.

### Patescibacteriota metagenome-assembled genomes

High-quality Patescibacteriota MAGs with completeness ≥ 90% and contamination <5%, as assessed with CheckM2 [39], were selected for further analysis of metabolic potential and taxonomy [38]. Phylogenetic tree was built with GTDBtk v. 2.7.2 de novo workflow (database 232) [40], using Rstudio packages ape v.5.8 [41], treeio v.1.22.0 [42] and ggtree v.3.6.2 [43]. The abundance of the Patescibacteriota was estimated with CoverM v.0.7.0 (methods tpm, covered_fraction) [44], and MAGs with horizontal coverage of ≥ 25% were considered present in a sample.

Metabolism of the MAGs was estimated with anvi’o v9 [45]. KEGG (v.116) KOs were annotated with anvi-run-kegg-kofams. Metabolic module completeness was estimated with anvi-estimate-metabolism, using pathwise completeness of ≥75% as threshold for classifying metabolic module as present in a MAG. The present metabolic modules were visualized in RStudio using the packages ggplot2 4.0.3 [46] and vegan 2.7-6 [47]. Metabolic similarities and dissimilarities of the Patescibacteriota MAGs were visualized using Principal Coordinates Analysis (PCoA), based on binary Jaccard distance of the presence/absence matrix of KO-groups. The host association of the MAGs was assessed with symclatron v0.10.10 [48].

### Viral metagenomic analysis

The modular viromics pipeline (MVP) v.1.0.0 [49] was used for viral detection and annotation, where viral contigs were predicted with geNomad v.1.7.4 (database 1.7) and their quality assessed with CheckV v.1.0.3 (database v1.5) [50]. Based on CheckV annotations, the geNomad predictions were retained only if the number of viral genes was ≥1, the ratio of host genes to viral genes was ≤1 and the sequence was ≥5kbp in length or ≥50% complete (by AAI prediction). Viral abundance across samples was calculated as reads per kilobase million (RPKM), and for further RPKM-based analyses, the 50% horizontal coverage threshold was used. The community dissimilarities were visualized with Non-metric multidimensional scaling (NMDS) utilizing Bray-Curtis distance and tested statistically with PERMANOVA (permutations = 9999, method = “jaccard”).

Host predictions for prokaryotic vOTUs were made with iPHoP v.1.4.1 [51] using the iPHoP_db_Jun25_rw database and a custom database constructed using all nonredundant MAGs obtained previously from the same dataset [38]. The predictions were filtered for the best matches only (in case of multiple host taxa predicted), and the outputs from two databases were combined at the level that they agreed. The cutoffs of 70 and 90 were used for family-and genus-level predictions, respectively. Viral lifestyle was predicted with Bacphlip v. 0.9.6 [52], and the vOTU lifestyle was considered as lysogenic if Bacphlip score was >0.5, MVP predicted it as provirus or the vOTU had PHROG annotation to integrase [53]. Otherwise, the vOTU was considered virulent. Statistical differences between the sample types and palsa frozen soil cores were tested with Kruskall-Wallis and Dunn’s test with Bonferroni correction, using log2 transformed ratio of temperate to virulent phages in RPKM%.

The vOTUs of ≥200 kbp in length and phages with a host prediction to Patescibacteriota were selected for additional genome annotations with Phold v.0.2.0 [54], and DRAM-v v. 0.1.2 (which was used at the KBase platform) [55]. To further validate predictions of interest, BLASTn and BLASTp searches were performed against core_nt and nr_cluster_seq databases, respectively (refs) using an E-value threshold of 1e-5, as well as HHPred searches against PDB_mmCIF70_20_Feb and SCOPe70_2,08 databases [56] with a threshold of 95% probability. The VIRIDIC webtool was used for analysing intergenomic distances between viral genomes [57]. tRNA genes in jumbo vOTUs and vOTUs linked to Patescibacteriota were predicted using ARAGORN v1.2.40 [58].

## Results

### Patescibacteriota were detected in the deepest palsa soils and small freshwaters

The metagenomics workflow yielded 57 medium and high-quality MAGs classified as Patescibacteriota, with an average completeness of 85.5% and contamination of 1.76%. The average genome size was 834 320bp (359 022 - 2 388 288bp), and CG content 47% (28-58%) (Table S1). The MAGs spanned six classes, 17 families and 23 genera. Eight MAGs could not be classified to the genus level. To explore the abundance and metabolism, only the high-quality MAGs with the completeness of ≥90% and contamination <5% [59], were chosen (n=33). These MAGs had slightly higher genome sizes (956 129bp) and lower CG contents (46%) than the Patescibacteriota MAGs overall. Most of the MAGs belonged to Patescibacteriota classes Minisyncoccia (17) and Saccharimonadia (11), and the rest to classes Doudnabacteria (2), Gracilibacteria (2), and Patescibacteriia (1) (Figure 1). Genus-level classification was missing for 6 MAGs: 2 Doudnabacteria (family VMFF01) and 4 Saccharimonadia (families DATKHC01 n=3 and UBA4665 n=1). Out of the high-quality MAGs, only 3 had an ANI value calculated and 30 MAGs had no ANI value, likely indicating a high novelty of these Patescibacteriota at the species level.

**Figure 1.**
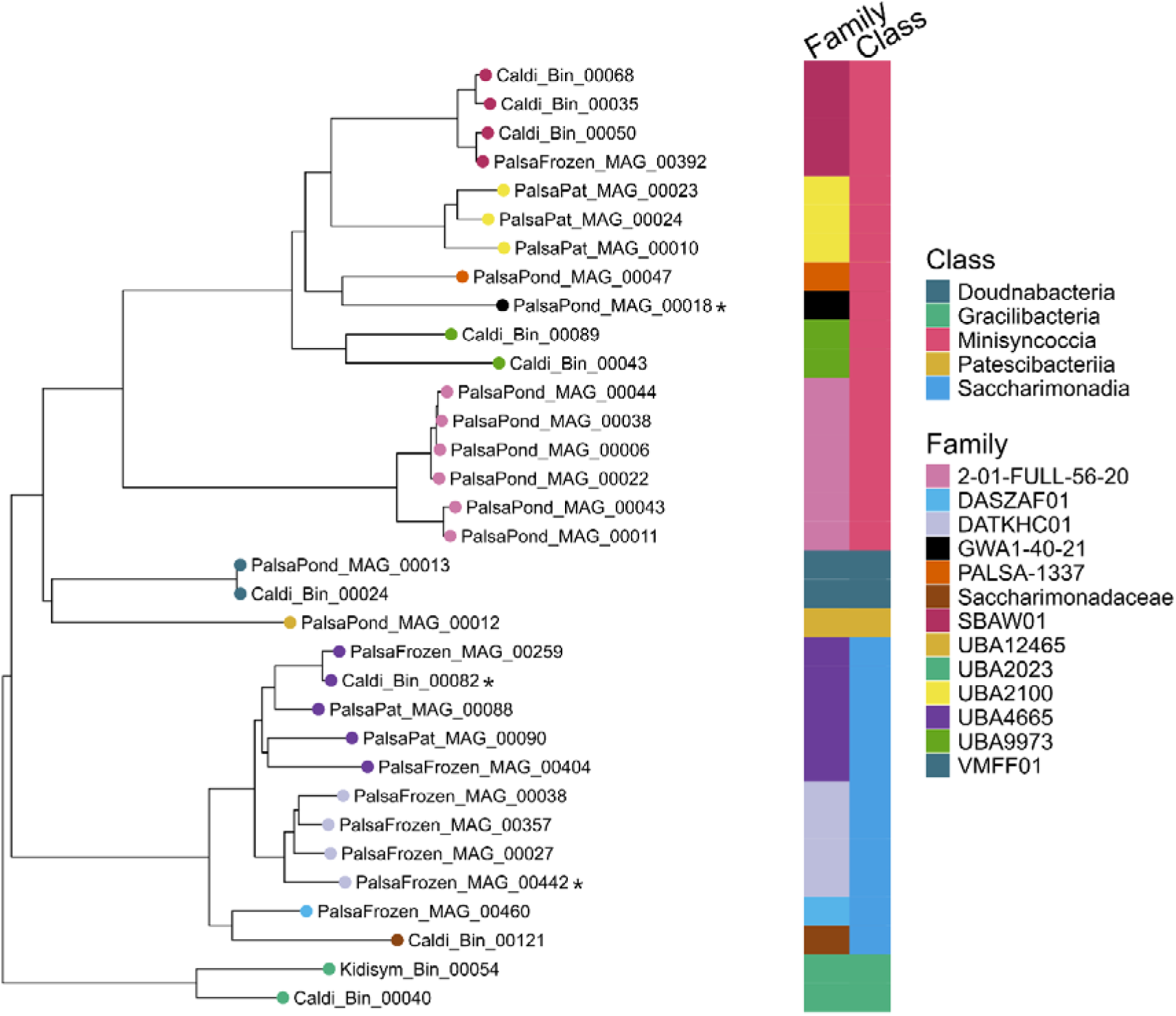
Phylogenetic tree of the high-quality metagenome assembled genomes (MAGs) of Patescibacteriota, with names indicating the assembly from where the MAG was retrieved. MAGs with names starting with PalsaPat or Caldi are retrieved from assemblies made from palsa soil core 26 deepest samples, Kidisym refers to assembly with stream water samples and PalsaPond to palsa ponds and PalsaFrozen to assembly from palsa frozen soil. Dots in front of the MAG name and in the first bar indicate family-level taxonomy, while the second bar indicates class-level taxonomy.

High-quality Patescibacteriota MAGs were most abundant in palsa waters, deeper layers of palsa permafrost and the stream sites. Notably, Patescibacteriota MAGs showed clear preference of terrestrial or aquatic habitats, with only three MAGs present in both types of environments (Figure 2). Saccharimonadia were present only in palsa soil, except for one MAG in the family Saccharimonadaceae, which was present in both palsa 26 deepest layers and palsa waters. Saccharimonadia were also present across all palsa soil profiles, while Minisyncoccia were only found in palsa soil profile 26. Minisyncoccia families were also divided between sample types, e.g. families SBAW01 and PALSA-1337 were only in palsa soil 26 deeper layers, whereas the family 2-01-FULL-56-20 was only found in palsa ponds (Figure 2).

**Figure 2.**
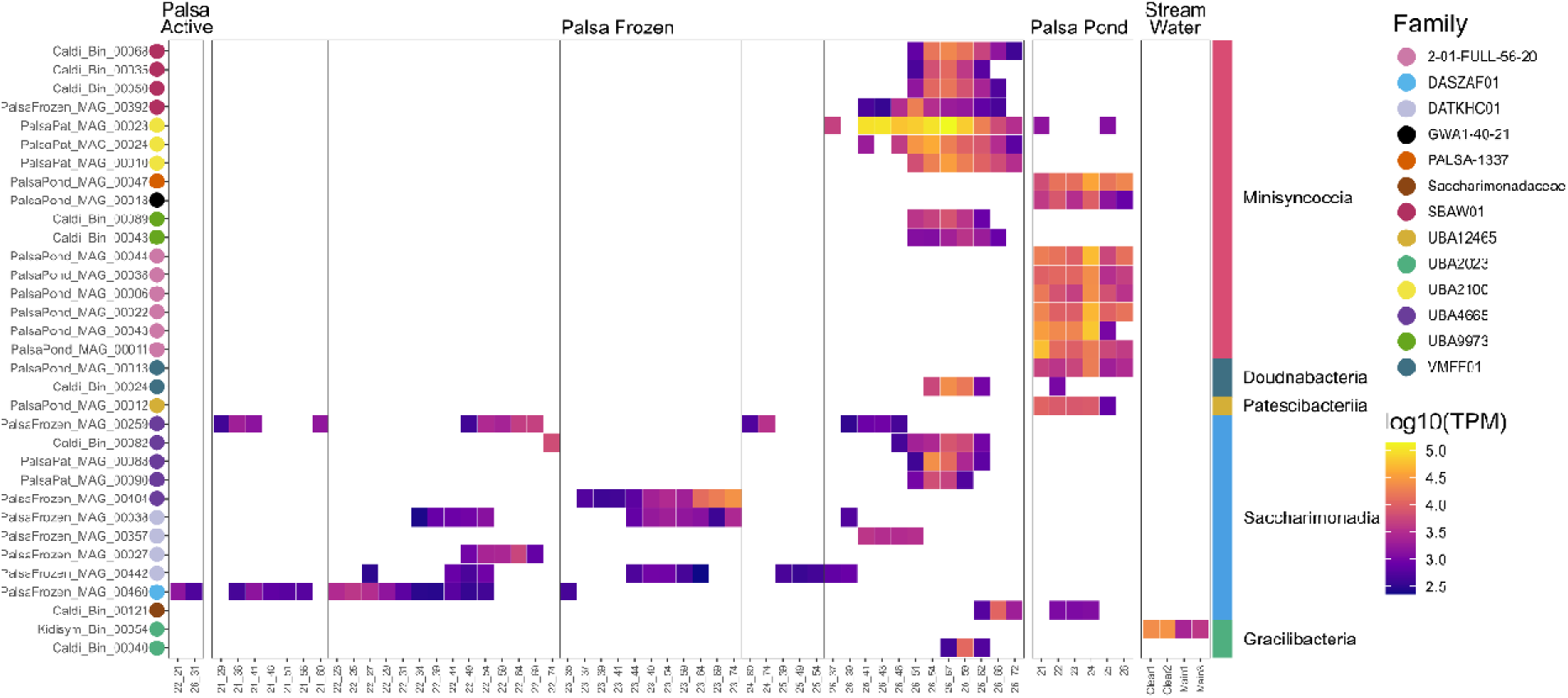
Abundance of high-quality Patescibacteriota MAGs in the subarctic palsa and peatland soils, ponds and streams normalized to log10(TPM). Only the samples with at least one high-quality Patescibacteriota MAGs are shown. The vertical lines divide palsa frozen soil profiles. The x-axis labels refer to sample code, and in palsa active and palsa frozen the first number refers to the sample and second number to the sampling depth. Family level taxonomic classification is shown as colour-coded dots after the MAG name on the left side of the heatmap, while class level taxonomy is marked on the right side.

### Patescibacteriota were host dependent, with most represented classes showing differences in metabolic potential

Patescibacteriota are characterized by their streamlined genomes, with highly reduced metabolic pathways indicative of symbiotic or parasitic lifestyle [20,35]. Our results were in line with this, since all of the high-quality MAGs were classified as host-associated (n=30) or obligate-intracellular (n=3) by symclatron. Host-associated lifestyle was also supported by all MAGs lacking complete TCA cycle, and a majority having incomplete glycolysis pathway and pentose phosphate pathway (Figure 3B, Table S2). Most of the MAGs also had incomplete glycolysis core 3 carbon pathway. Almost all the MAGs encoded genes for complete F-type ATPase (M00157). The lack of electron transport chain complexes, but presence of F-Type ATPase has been detected across multiple Patescibacteriota [10].

**Figure 3.**
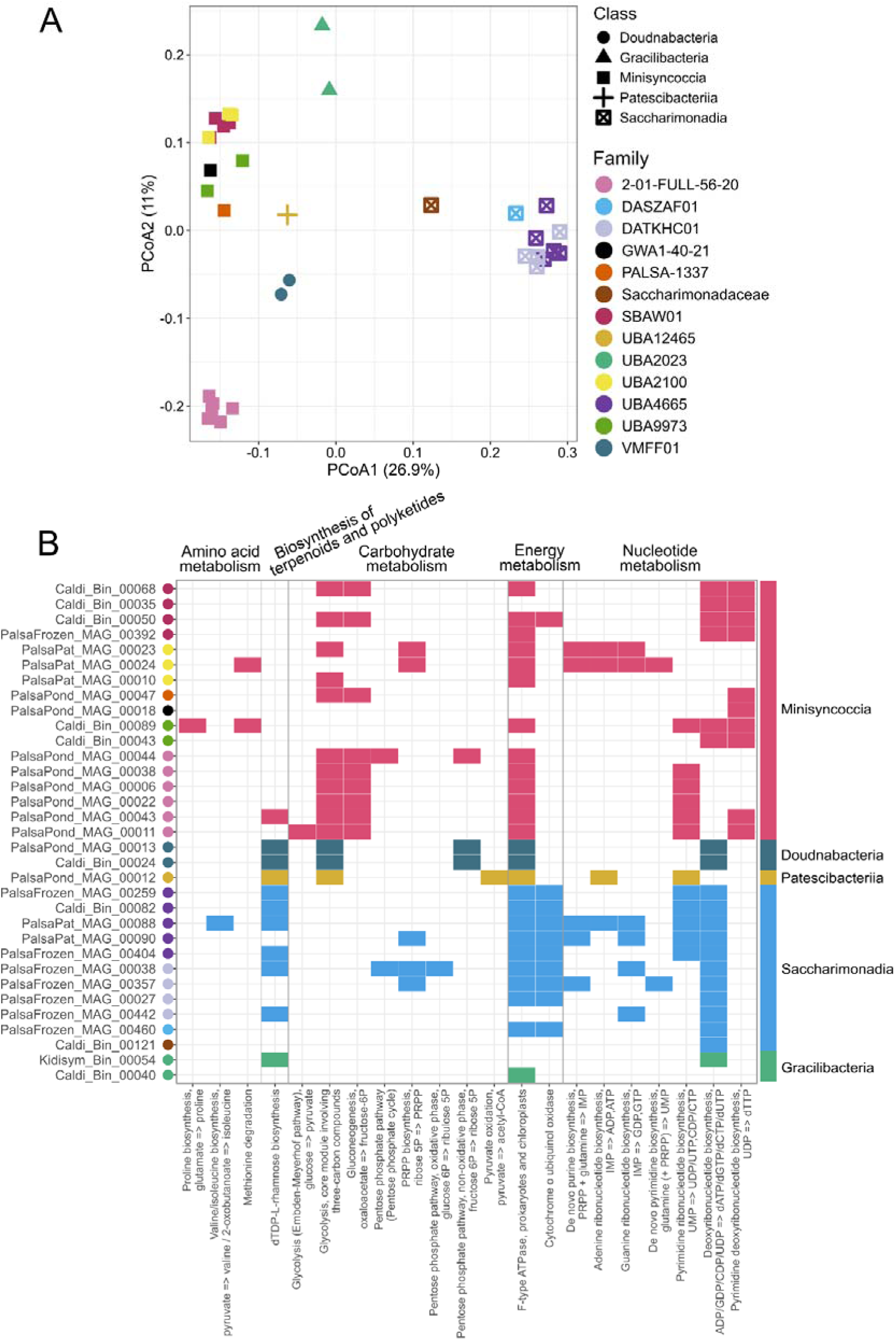
Principal Coordinates Analysis (PCoA) with binary Jaccard Distance of all KEGG K0-groups in high-quality Patescibacteriota (A). Shape refers to the class-level taxonomy and color to the family-level taxonomy. All complete metabolic pathways and modules of the high-quality Patescibacteriota MAGs with threshold of 75% pathwise completeness are considered complete (B). Color after MAG name indicates family-level classification with colors matching the PCoA, and class-level taxonomy is indicated on the right side of the figure.

Clear differences in the metabolic potential between Patescibacteriota classes were observed, especially for the most numerous classes of Minisyncoccia and Saccharimonadia (Figure 3A). Almost all Saccharimonadia MAGs had dTCP rhamnose biosynthesis, indicating rhamnose as cell wall material and possibly rods in cell wall [60], whereas only one Minisyncoccia MAG had this module complete. Saccharimonadia also had cytochrome o ubiquinol oxidase pathway, which again was missing in almost all Minisyncoccia MAGs. This pathway had been proposed as a possible protection method for obligatory anaerobes against oxygen [21], or as a pathway for fermentative microbes to occasionally use aerobic respiration [18]. The majority of Minisynccoccia MAGs had the three-carbon core glycolysis pathway and gluconeogenesis pathways. Gluconeogenesis is responsible for utilization of acetate in oligotrophic environments [61].

Patescibacteriota metabolic potential has been described from subglacial environments [62] and we compared MAG KO-groups between the subglacial and our permafrost ecosystems. Of the KOs detected in subglacial environments, assembly proteins for type IV pilus were widely present in MAGs (pilD K02654 (33), pilM K02662 (30), pilT K02669 (26)). Notably, Minisyncoccia had PilO pilus assembly protein (14 MAGs), whereas Saccharimonadia lacked it completely. Type IV pilus is responsible for twitching motility of the cells, DNA uptake and cell adherence [63]. Competence proteins ComEC (K02238) and ComFC (K02242) were present in all MAGs, except Saccharimonadia. All Patescibacteriota had superoxidase dismutase, which is a mechanism to protect the cells against reactive oxygen. However, Saccharimonadia MAGs had nickel superoxide dismutase (K00518), whereas Minisyncoccia and the rest of the MAGs had Fe-Mn family superoxide dismutase (K04564). Antitoxin system TA II (K07272) was present in one third of the MAGs (20). Almost all the MAGs (29) had the gene for peptide/nickel transport system substrate binding protein (K02035) (Table S3).

All MAGs lacked ribosomal protein L30 (K02907). Minisyncoccia MAGs lacked L1 ribosomal protein (K02863), which was found in all other MAGs. All Saccharimonadia lacked L29 (K02904) and L32 (K02911), but these were also missing in most of the Minisyncoccia. All MAGs of the family 2-01-FULL-56-20 (Minisyncoccia) lacked ribosomal protein L28 but had L29 unlike the rest of the Minisyncoccia families. Family SBAW01 (Minisyncoccia) lacked S21. All MAGs lacked CRISPR-related proteins (K02048). CRISPR systems have shown to be reduced from Patescibacteriota genomes [35,64] with certain exceptions [33].

### Viral community composition and lifestyle was determined by the sample type

The metagenomic samples yielded 16 354 viral operational taxonomic units (vOTUs), of which 436 were high-quality or complete (Table S4). The majority of vOTUs represented the class *Caudoviricetes* (92.6%, 15 138), i.e., dsDNA tailed prokaryotic viruses. About 3% of all vOTUs (492) were classified as giant viruses from the class *Megaviricetes,* while the rest were mainly from classes *Maveriviricetes, Tectiliviricetes* and *Polintoviricetes* (Figure S1). Of all the vOTUs, 3.3% (541) were identified as proviruses.

The length of vOTUs varied from 1 679 to 530 549 bp, with a mean length of 13 532 bp (Figure S2 & Table S4). Seventeen vOTUs had genomes of over 200 kbp in length, including 16 tailed prokaryotic viruses classified as *Caudoviricetes* and one giant virus from the class *Megaviricetes*. Thus, these 16 *Caudoviricetes* vOTUs represented jumbo phages [65,66]. Moreover, one vOTU was exceptionally large, 530 kbp in length, exceeding the 500-kbp threshold defined for megaphages [67]. This megaphage was detected only in palsa water samples. According to VIRIDIC, all the jumbo phages represented different genera. Notably, as predicted by Phold, four and nine jumbo phages encoded ribosomal protein S21 (a structural component of the small ribosomal subunit) and ribosomal protein S6 glutaminyl transferase (post-translational modification of ribosomal components), respectively (Tables S5-S6). DRAM-v annotation also suggested a large subunit ribosomal protein L7/L12 in addition to S21 in one of the jumbo phages. In addition, multiple tRNA genes were predicted in most jumbo phages: 21-56 genes in all jumbo phages, except that two had only one tRNA gene and four had no such genes predicted (Table S7). In total, using Phold, 655 CDSs in 16 jumbo phages were annotated as “moron, auxiliary metabolic gene and host takeover”, including putative glycosyltransferases, ABC transporters, virulence factors, sporulation proteins, toxins, resistance, anti-CRISPR proteins and a CRISPR/Cas-associated protein (Tables S5-S6). However, reliable functional assignments of these genes remain challenging and would need further validation.

NMDS of the vOTUs divided the samples by the X-axis into two groups: palsa soils and palsa ponds; and peat soils, peat ponds and stream samples (Figure 4A). Notably, some of the palsa soil communities were grouped closer to the palsa ponds. These distinct layers were the deepest sampled layers (depth >40cm) from the palsa soil profile 26. The highest number of unique and overall vOTUs were discovered from the palsa waters, which had 6 681 unique vOTUs present, followed by 3 095 vOTUs in palsa frozen soils and 1 593 vOTUs in stream waters (Figure 4B). Notably, there was only one vOTU detected across all sample types and overall, a fairly low number of vOTUs were present across multiple sample types. PERMANOVA analyses revealed that soil depth, sample type, pH, SOM, and C:N ratio each significantly contributed to differences in viral community composition among samples (R² of 22.91, 9.76, 2.89, 1.80, 1.34, respectively, p < 0.01, all parameters tested separately). Thus, soil depth and sample type explained the greatest proportion of variation in the viral community composition.

**Figure 4.**
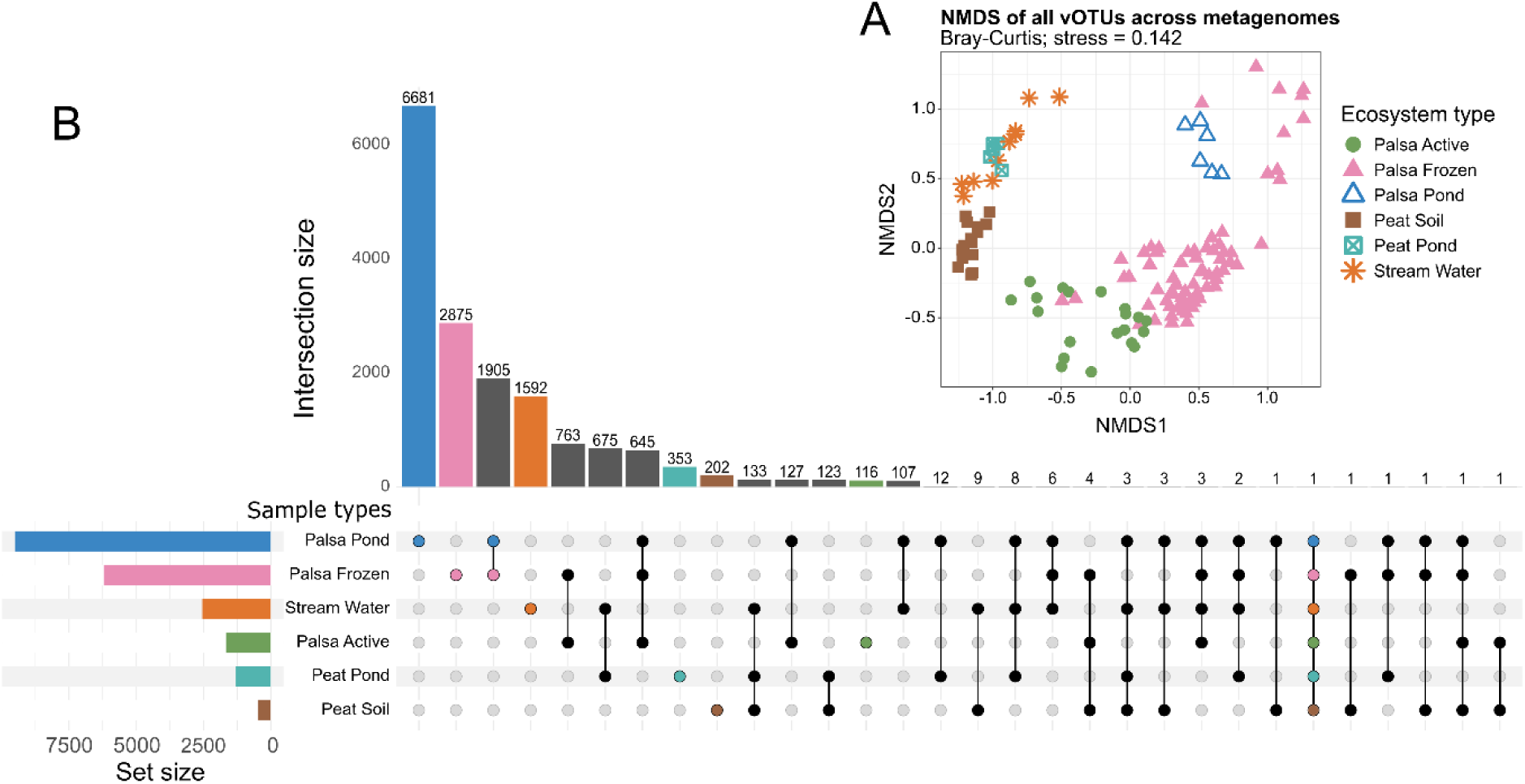
Non-metric multidimensional scaling (NMDS with jaccard binary distance) of all viral operational taxonomic units (vOTUs) in frozen and active palsa soil, palsa ponds, peat soils, peat ponds and stream water samples (A). The abundance of vOTUs was normalized to reads per kilobase million (RPKM). The presence of shared and unique vOTUs across sample types, with RPKM >0 & HC >0.5 considered present (B).

Approximately 40% of all vOTUs (6 488) were *in silico* linked to their putative prokaryotic hosts: 38.5% (6 299) and 1.2% (189) to Bacteria and Archaea, respectively. Of all these predictions, 32% (2 103) and 21% (1 380) were based on custom and default iPHoP databases respectively, while 46% (3 044) were retrieved from both. The host predictions spanned a wide range of taxa: 43 phyla, 109 classes, 598 families and 331 genera. The most commonly predicted host phyla were Pseudomonadota (2 289, Acidobacteriota (1 036), Bacteroidota (685), Actinomycetota (425) and Bacillota (648) (Figure 5A). The most numerous host class-level predictions included Alphaproteobacteria (1169), Gammaproteobacteria (1094), Terriglobia (1 023) and Bacteroidia (678). From the archaea, Halobacterota (50), Thermoproteota (45), Nanobdellota (24), Asgardarchaeota (20) and Methanobacteriota (18) were the most predicted phyla. About 66% (355) of proviruses were linked to a putative host at the phylum level, including Pseudomonadota (122), Acidobacteriota (117), Actinomycetota (35), Bacillota (20) and Bacteroidota (17). Pseudomonadota and Acidobacteriota phages were the most abundant in palsa soils, palsa ponds and peat soils (Figure 5A-B). Pseudomondota phages had a similar pattern of relative abundances across all samples. Actinomycetota- and Chloroflexota-linked phages were more abundant in palsa frozen soils than the rest of the sites. Planctomycetota phages were more abundant in the active palsa soil layers than frozen.

**Figure 5.**
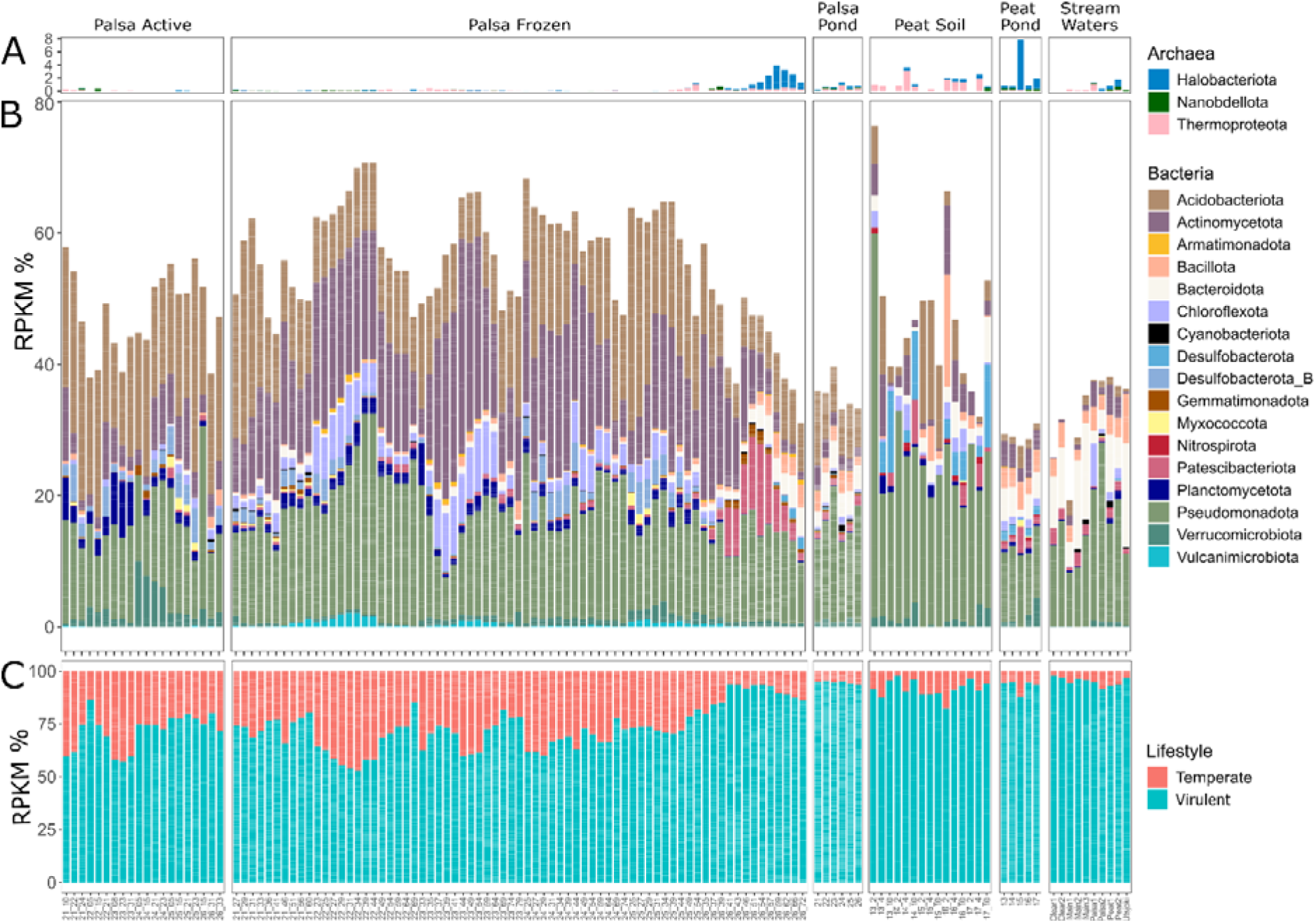
Relative abundance of the prokaryotic viruses grouped by their predicted hosts on phylum level for archaea (A) and bacteria (B). Abundance of virulent and temperate prokaryotic viruses across the samples (C). Labelling in the x-axis refers to sample name, and with soil samples the sample name is followed by sampling depth in centimeters.

With the Bacphlip lifestyle prediction score threshold of 0.95, 24% (3 842) of prokaryotic viruses were determined as virulent and 4% (637) as temperate, while the rest were unclassified (Figure S3). With the combined approach (with lower threshold and additional data from MVP annotations, see Methods), 90% (14 286) of the prokaryotic vOTUs were identified as virulent and 9.6% (1 520) as temperate. Temperate phages were more abundant in palsa active and palsa frozen soils compared to the peat soils, peat and palsa ponds and streams (Kruskall-Wallis test, followed by Dunn’s test with Bonferroni correction; p<0.05, Fig 5C). Notably, in palsa frozen the soil profile 26, the deeper layers had overall lower relative abundance of temperate phages than in the other palsa soil profiles. The palsa active soil profile 26 differed also statistically from the frozen soil profiles 22, 23 and 24 (Kruskall-Wallis test, followed by Dunn’s test with Bonferroni correction; p<0.05), but not from the profiles 21 and 25 (p=0.09 and p=0.08, respectively).

### Patescibacteriota phages represent diverse genera

From the viral analysis, 196 vOTUs had Patescibacteriota as their predicted hosts, of which 4% (n=8) were complete or high-quality vOTUs (Table S4). The host prediction was made based on the custom iPHoP database for 55% of the vOTUs (107), 27% by iPHoP default database (53), and 18% by both databases (36). The genome length in Patescibacteriota-associated vOTUs varied from 5 015 to 170 127 bp, with a mean length of 14 539 bp. The majority of these vOTUs were putatively classified within the class *Caudoviricetes* (189), of which ten were classified also at the family level as *Straboviridae* (4), *Kyanoviridae* (2), *Ackermannviridae* (1), *Mesyanzhinovviridae* (1), *Schitoviridae* (1) or *Vilmaviridae* (1). One vOTU was placed within the class *Tokiviricetes*, family *Lipothrixviridae,* which is most likely a misclassification, since this family comprises rod-shaped viruses that infect hyperthermophilic archaea [68]. Five vOTUs remained unclassified, and one was classified only at the kingdom level as *Bamfordvirae*. The latter was predicted to encode tail proteins by Phold, so its classification could not be reliably resolved. Based on the VIRIDIC results, only two vOTUs belonged to the same genus (PalsaFrozen_k157_2560995 and PalsaPate_k157_358477), indicating high taxonomic diversity among the Patescibacteriota-linked phages. Out of the 196 vOTUs, 4% (8) were proviruses, which were all classified as *Caudoviricetes*.

The Patescibacteriota vOTUs targeted hosts mainly from the classes Minisyncoccia (125, 5 proviruses), Saccharimonadia (47, 1 provirus), Microgenomatia (8), and Patescibacteriia (5, 1 provirus). The six proviruses of Minisyncoccia had host predictions to the order UBA9973. Eleven vOTUs had predictions for both families of UBA21200 (Minisyncoccia) and RAAP-2 (Actinomycetota) with custom databases. None of the prediction methods utilized CRISPR spacers, supporting the observation that most Patescibacteriota lack CRISPR systems due to genome streamlining [13,35]. Nonetheless, one of the Patescibacteriota-associated vOTUs (PalsaCaldi_k157_412198) had an anti-CRISPR system protein annotated with Phold (see below for Phold annotations), suggesting that CRISPR-systems are still relevant in antiviral defense within this group. Indeed, the presence of CRISPR-Cas systems has been previously shown for some Patescibacteriota lineages [34].

Patescibacteriota vOTUs were most abundant in the palsa soil profile 26 deepest layers (depth >40cm), which was largely aligned with the abundance of the Patescibacteriota MAG abundance (Figure 6A-B). In some of the sampled soil layers, up to 7-10% of the observed prokaryotic viruses were those linked to Patescibacteriota. Bacphlip (with the 0.95 threshold used) predicted 47 viruses as virulent and 2 as temperate. The combined prediction strategy (see above and Methods) suggested 182 phages to be virulent and 14 to be temperate. Notably with both methods, virulent phages were more abundant in most of the samples (Fig. 6C).

**Figure 6.**
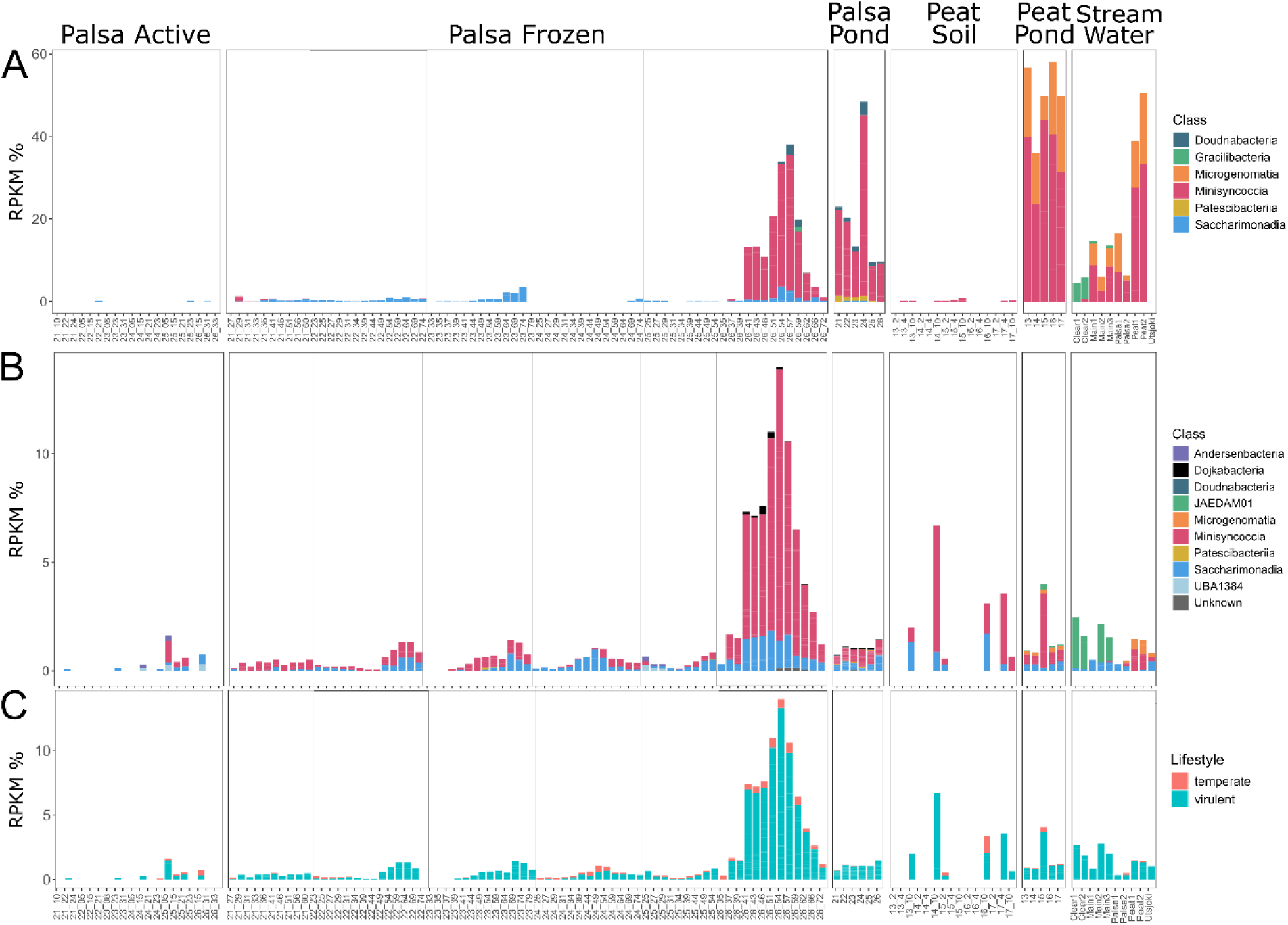
Relative abundance of medium- and high-quality Patescibacteriota MAGs normalized to RPKM% (A). Relative abundance of the Patescibacteriota phages normalized to RPKM% (B). Please note that the class marked as JAEDAM01 corresponds to the class Gracilibacteria in A. Relative abundance of Patescibacteriota phages by lifestyle prediction (C). Labelling in the x-axis refers to sample name, and with soil samples the sample name is followed by sampling depth in centimeters.

### Viral Genome Annotations of Patescibacteriota-associated vOTUs

The genome annotations of Patescibacteriota-linked vOTUs yielded 4 223 coding sequences (CDS) in total (predicted by Phold) and varying numbers of tRNA genes per vOTU: most of these vOTUs lacked tRNA genes (176 out of 196), 13 vOTUs had 1-3 such genes, and seven vOTUs had 7-30 tRNA genes (Table S7). The largest genome (170 kbp) had the highest number of tRNA genes predicted (30). Most of the CDS had unknown functions (2 407) or were categorized as other (294) by Phold. The assigned categories included DNA, RNA and nucleotide metabolism (608), head and packaging (329) or moron, AMG or host takeover (128). Two vOTUs had genes putatively encoding small subunit ribosomal protein S12, which was predicted both by Phold and DRAM-v (Tables S9-S10). Other predictions related to modulation of host translation included proteins involved in tRNA metabolism and translation-related processes, e.g., components of the queuosine biosynthesis pathway, tRNA processing enzymes, and translation-associated proteins. Two vOTUs also encoded chaperonin *groES* (K04078, Table S10). As predicted both by Phold and DRAM-v, several vOTUs encoded proteins that could potentially enhance nucleotide metabolism, such as thymidylate synthases.

Putative predictions of proteins involved in virus-host interactions included DarB-like antirestriction proteins, virulence factor proteins, and a putative Zot-like toxin. The latter one retrieved hit to zinc proteases in HHPred searches (>99% probability hits) and BLASTp hits to insulinase family proteins in Patescibacteriota, leaving a precise function of this protein uncertain. In addition, a protein associated with CRISPR/Cas system and two Cas4-domain exonucleases were predicted, which were also verified by HHPred searches (>99% probability hits to Cas4 exonucleases retrieved). However, it remains unclear whether these proteins could take a role in interactions with the host CRISPR/Cas systems or have been co-opted for more general DNA processing functions. Interestingly, Patescibacteriota phages have been previously reported to encode possible AMGs related to Patescibacteriota cell modification to attach to their prokaryotic symbionts [35]. According to DRAM-v predictions, ten Patescibacteriota phages encoded GDP-mannose-dehydratases (Table S10), which could be potentially involved in O-antigen nucleotide sugar biosynthesis and thus contribute to the surface adhesion and cell-cell aggregation of Patescibacteriota and other microbial cells [35]. Experimental validation would be needed to verify this speculation.

## Discussion

### Minisyncoccia and Saccharimonadia showed clear patterns of ecosystem preference and metabolic capacity

Our aim was to characterize the Patescibacteriota and their role in the microbial communities between sample types in a sporadic permafrost ecosystem. The retrieved MAGs were mainly from the classes Saccharimonadia or Minisyncoccia, which are generally the most abundant classes of Patescibacteriota across ecosystems [69]. Consistently with other reports [28,69–71], Patescibacteriota MAGs recovered in this study were relatively short in length and had highly incomplete metabolic pathways, indicating a host-dependent lifestyle. Host-dependent lifestyles were also confirmed with predictions based on symbiotic genomic signatures. All Patescibacteriota MAGs encoded genes related to motility and uptake of external DNA to supplement their incomplete metabolism and reduced genomes, which has been also described in cultivation studies [24].

Patescibacteriota exhibited a strict presence and absence pattern by sample type. Saccharimonadia was present in the likely oxic and frozen soil layers. Saccharimonadia has been previously shown to have stronger preference for oxic conditions in groundwater samples than other Patescibacteriota classes [72]. However, Patescibacteriota may switch metabolic processes to suit both oxic and anoxic conditions [72]. Minisyncoccia was only present in palsa thaw ponds and the likely anoxic deeper layers of degrading palsa soil. The previously isolated Minisynccoccia strain supports our finding of Minisyncoccia being mainly detected in likely anaerobic samples, as the class was previously classified as facultatively anaerobic [24]. Notably, the palsa thaw ponds were oxic [37] but as other Patescibacteriota, the Minisyncoccia classes present in the palsa ponds could also be adapted to oxygen [72].

Another factor in the Patescibacteriota abundance is the presence of their hosts, especially as all of the MAGs of this study were classified as host-associated. Minisyncoccia has been previously isolated in association with methanogenic archaea Methanothrix [24]. Minisyncoccia in our study were recovered from soil layers with high abundance of archaea annotated as methanogenic Bog-38 (Halobacterota) [38], suggesting that it could potentially act as the host. The cultivated strain of Minisyncoccia slowed both the growth and methane production rates of the methanogenic host [24].

### Viral community composition shows distinction by sample types

Predicting viral sequences in the bulk metagenomes of permafrost and peat soils and adjacent small-scale freshwaters has yielded a high number of vOTUs (16.4 K), most of which represented tailed dsDNA viruses of bacteria of archaea (*Caudoviricetes*). Viruses belonging to the class *Caudoviricetes* are among the most recalled groups of viruses predicted from meta-omics datasets in various environments, including soil and freshwater ecosystems [73,74]. The vOTUs had very little overlap across sample types, which has been observed across studies on soil viromes [30,75].

Tailed phages recovered in this study were *in silico* linked to bacterial and archaeal groups that are involved in important ecological processes, such as methanogenesis. Their distribution across sites has mostly mirrored patterns observed for their host communities [38], suggesting coupled virus-host dynamics. Interestingly, temperate phages were overall more abundant in palsa soils than peat soils and all water samples. However, most of the phages were predicted as virulent. Although bulk soil metagenomes likely miss a substantial diversity of viruses present as extracellular viral particles, our results are still comparable across soil samples, as they were processed following the same protocols. Virulent lifestyle is generally more common in permafrost soils, and soils overall [30,75,76]. However, a higher prevalence of temperate phages in palsa soils compared to peat soils in this study may reflect local environmental conditions (lower pH, higher soil organic matter and total C to total N ratio [37]) and host community characteristics that favor lysogeny. Longer monitoring efforts would be needed to verify these trends and their underlying factors.

The recovery of 16 jumbophages, including one megaphage with the genome length of 530 kbp added further insights into virus-host interactions in permafrost-affected environments. Consistent with other reports, these phages demonstrated an expanded genetic repertoire with multiple tRNA genes, supporting a possible extensive interception of host biology [77]. In addition, the detection of vOTUs representing other groups of viruses, i.e. giant viruses (class *Megaviricetes*), virophages (class *Maveriviricetes*, recently renamed to *Virophaviricetes*, [78,79] and polintons (class *Polintoviricetes*), highlighted eukaryote-associated viruses and virus-like elements in Arctic permafrost-affected ecosystems.

Patescibacteriota were among the most abundant viral hosts, indicating that they are likely an important component of the microbial and viral communities in the degrading permafrost and small freshwaters. Patescibacteriota phages are highly novel, with only few reports based on metagenomic data [28,34–36]. We discovered that Patescibacteriota phages were diverse, with almost all the 196 vOTUs belonging to different genera. Patescibacteriota phage dynamics seemed to be coupled with their hosts in the frozen palsa soils, as the phage and host abundance followed mostly the same pattern. The majority of the Patescibacteriota phages were predicted to be virulent rather than temperate. This is in contrast to previous findings from groundwaters, where Patescibacteriota phages were suggested to be mainly temperate [35]. Notably, experimental evidence for Patescibacteriota-phage interactions remains extremely scarce. The only cultivated model system described to date showed that phage infection dynamics can depend on host growth mode, spatial organization, and interactions between the Patescibacteriota epibiont and its bacterial host, resulting in either lysogenic or lytic outcomes under different conditions [36]. Additional cultivation-based studies and direct virus-host validation are needed to resolve how phages influence Patescibacteriota ecology and their complex interactions with host organisms in natural ecosystems.

To conclude, our results revealed the Patescibacteriota and their phages to be one of the most abundant groups in microbial and viral communities of a permafrost ecosystem. The most abundant Patescibacteriota classes, Minisyncoccia and Saccharimonadia, showed clear differentiation by their abundance across sample types and by metabolic potential of the cells. Patescibacteriota phages mostly followed the relative abundance patterns of their hosts. The potential impact of the Patescibacteriota on its host should be examined further, as Patescibacteriota was one of the most abundant phyla in the degrading permafrost soils. Currently, the dynamics between Patescibacteriota and its host remain scarce due to a low number of isolates. There is a critical need to improve understanding of this enigmatic phylum, and the impact on the hosts cell metabolism, especially through cultivation studies and in environmental samples. In addition, our results highlight the need for further research on the complex dynamics between Patescibacteriota phages, Patescibacteriota and their host cells with both metagenomics and culture-based methods, especially in the northern permafrost ecosystems facing rapid degradation.

## Data availability

Sequencing data is available in ENA projects PRJEB121465 and PRJEB85624. MAGs and vOTUs are accessible in Figshare: https://doi.org/10.6084/m9.figshare.34898898 (MAGs) and https://doi.org/10.6084/m9.figshare.34896960 (vOTUs)

## Acknowledgements

We thank Kevo Subarctic Research Station (University of Turku), Samu Elovaara and Janina Kariluoto for assistance with the field campaign. We thank CSC—IT Center for Science for the computational resources. We acknowledge DNA Sequencing and Genomics Laboratory (supported by HiLIFE and Biocenter Finland funding), Institute of Biotechnology, University of Helsinki for sequencing.

## CRediT statement

Nea Tuomela (Conceptualization, Funding acquisition, Field work, Data curation, Formal analysis, Investigation, Visualization, Writing—original draft, Writing—review & editing), Tatiana Demina (Data curation, Formal analysis, Writing—original draft, Supervision, Writing – review & editing), David N. Thomas (Conceptualization, Project administration, Field work, Investigation, Resources, Supervision, Writing— review & editing), Hermanni Kaartokallio (Conceptualization, Supervision, Investigation, Resources, Writing—review & editing), and Jenni Hultman (Conceptualization, Supervision, Investigation, Resources, Writing—review & editing).

