## Supplementary Figures for "Ultra-small bacteria and their phages discovered through metagenomics in a sub-arctic degrading permafrost ecosystem"

Authors & affiliations:

Nea Tuomela^1*^, Tatiana Demina^2,4^, David N. Thomas^1^, Hermanni Kaartokallio^3^, and Jenni Hultman^4^

^1^ Department of Environmental Sciences, Faculty of Biological and Environmental Sciences, University of Helsinki, Helsinki, Finland

^2^ Department of Microbiology, Faculty of Agriculture and Forestry, University of Helsinki, Helsinki, Finland’

^3^ Marine and Freshwater Unit, Finnish Environment Institute, Helsinki, Finland

^4^ Soil Ecosystems, Natural Resources Institute Finland, Helsinki, Finland

*Corresponding author Nea Tuomela, Department of Environmental Sciences, Faculty of Biological and Environmental Sciences University of Helsinki, Viikinkaari 1, 00790 Helsinki,

### Supplementary Figures

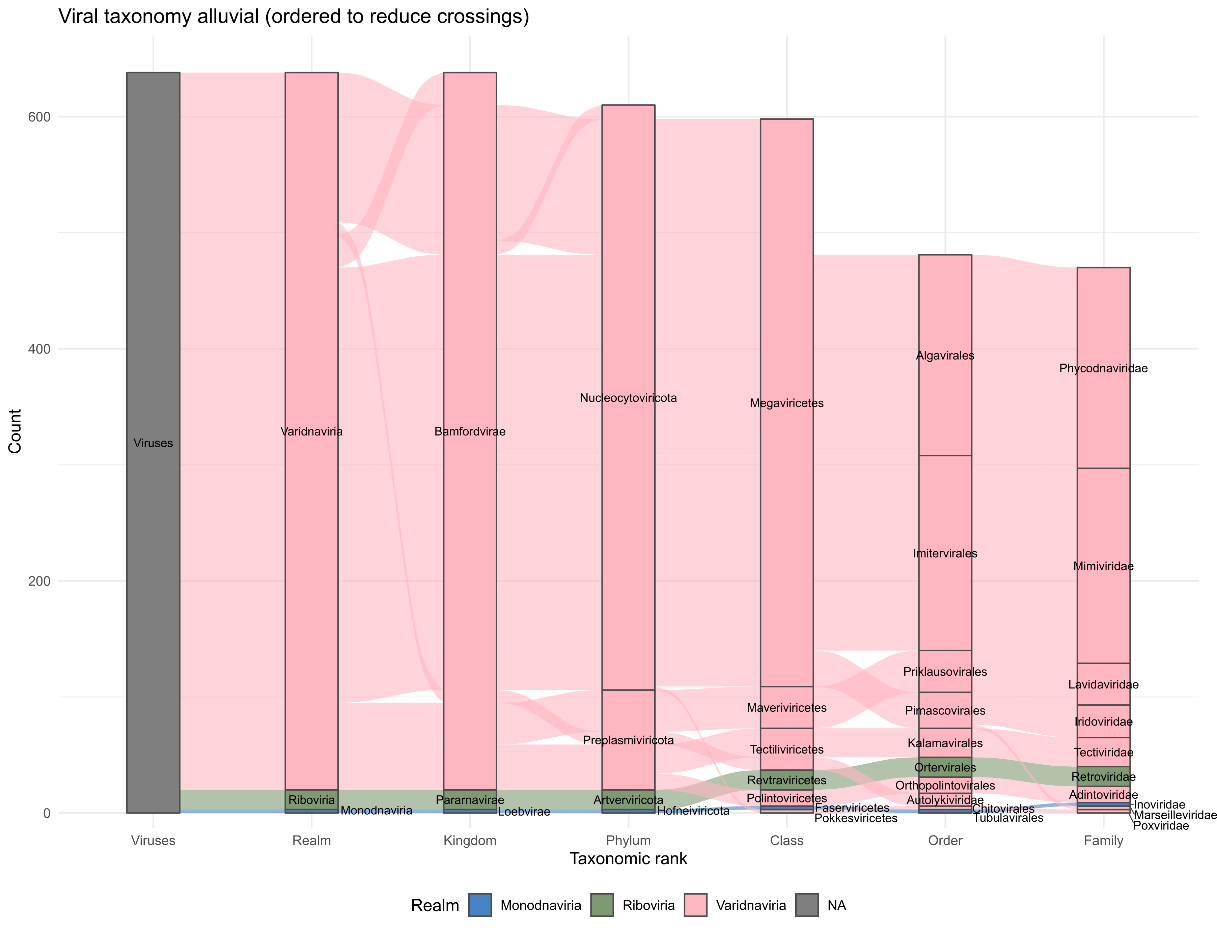

Supplementary Figure 1. The number of viral operational taxonomic units (vOTUs) per taxonomic rank, not including *Caudoviricetes* or unclassified vOTUs. Please note that the class *Maveriviricetes* was renamed to *Virophaviricetes* (Koonin et al., 2024; Simmonds et al., 2026).

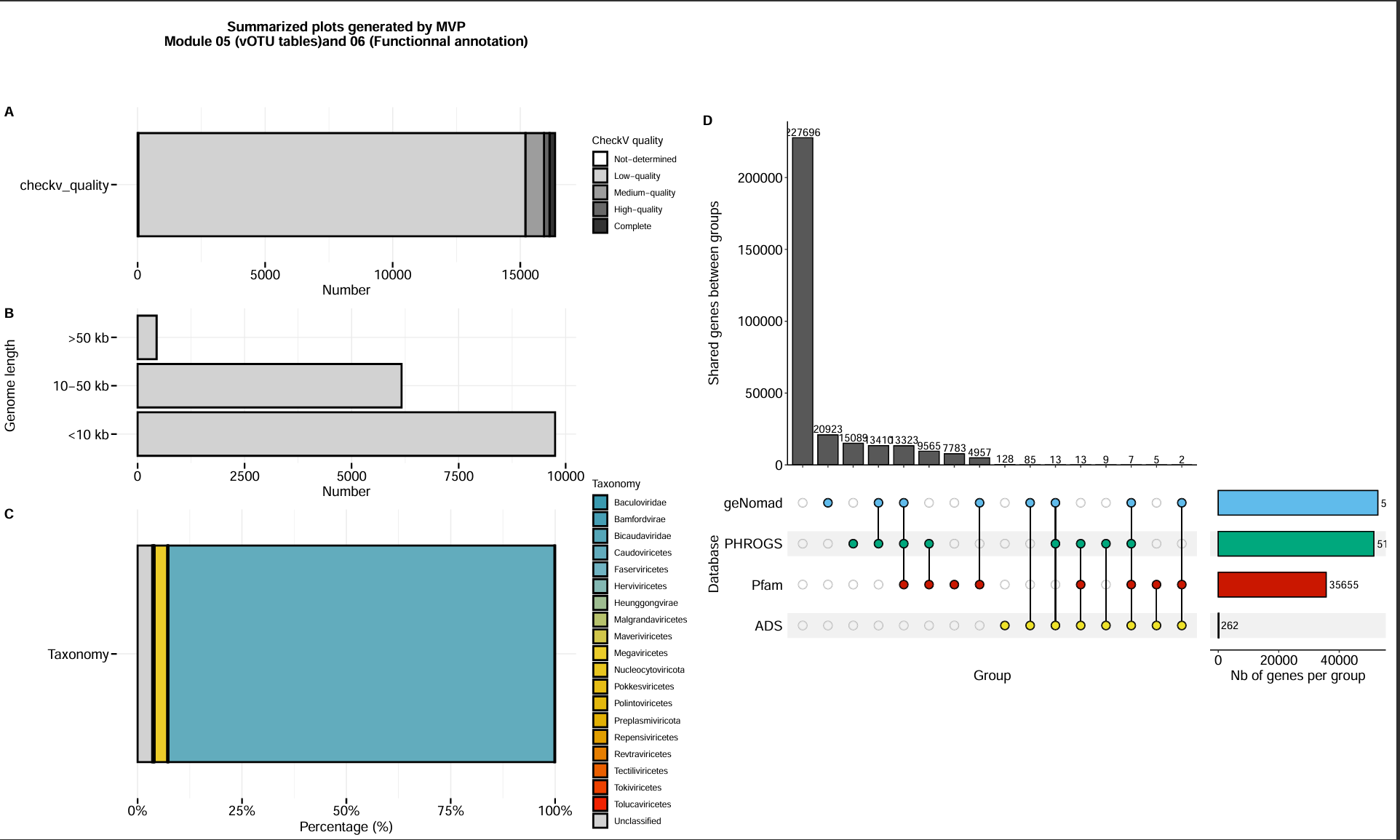

Supplementary Figure 2. The quality (A), genome length (B), taxonomy (C) and shared gene annotations (D), summarising the results of the modular viromics pipeline (MVP).

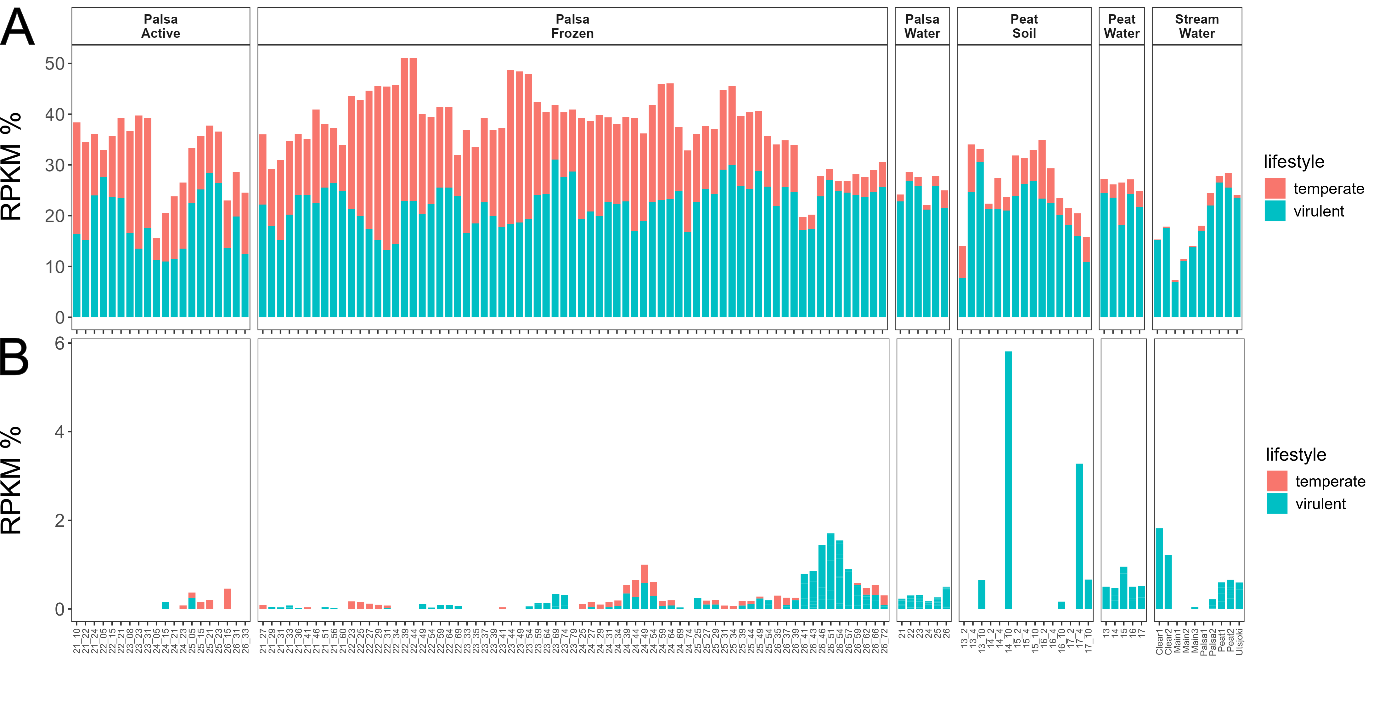

Supplementary Figure 3. Viral lifestyle predictions of all (A) and Patescibacteriota vOTUs (B) predicted with Bacphlip results with a threshold of 0.95.
